# Mental Tasks Induce Common Modulations of Oscillations in Cortex and Spinal Cord

**DOI:** 10.1101/2024.11.08.615786

**Authors:** Patrick Ofner, Dario Farina, Carsten Mehring

**Affiliations:** Bernstein Center Freiburg & Faculty of Biology, University of Freiburg, Freiburg im Breisgau 79104, Germany; Department of Bioengineering, Faculty of Engineering, Imperial College London, London W12 0BZ, United Kingdom; Bernstein Center Freiburg & Faculty of Biology, BrainLinks-BrainTools, University of Freiburg, Freiburg im Breisgau 79104, Germany

## Abstract

We investigated whether power modulations of cortical oscillations induced by mental tasks are paralleled by the same modulations in spinal motor neurons. We recruited 15 human participants and recorded high-density electromyography signals (HD-EMG) from the tibialis anterior muscle, as well as electroencephalography (EEG) signals. The cumulative spike train (CST) was computed from the activity of spinal motor neurons decoded from HD-EMG signals. The participants performed sustained dorsiflexion concurrent with foot motor imagery, hand motor imagery, mental arithmetic, or no specific mental task. We found significant power correlations between CST and EEG across trials irrespective of the mental task and across mental tasks at the intra-muscular coherence peak (Kendall’s *τ* coefficient *τ*_*trial*_ = 0.08*±*0.10, *τ*_*task*_ = 0.33*±*0.19, respectively; mean*±*std. dev.). CST power in beta and low-gamma bands could provide a novel control signal for neural interface applications, as power changes in these bands are not translated into actual force changes. To evaluate the potential of CST bands as a control signal, we classified the mental tasks from CST bandpower with a linear classifier and obtained classification accuracies slightly but significantly above chance level (30%±5%; chance level = 25%). These results show that mental tasks can modulate the power of cortical and spinal oscillations concurrently. This supports the notion that movement-unrelated oscillations can leak down from the cortex to the spinal level.

**Impact Statement:** Spike trains of spinal motor neurons exhibit frequency components above 10 Hz, which may partly reflect force-unrelated cortical oscillations and are modulated by mental tasks.

## 1 Introduction

Spinal motor neurons (MNs) within the same motor pool receive common synaptic input (Farina and Negro, 2015). This common synaptic input can be revealed by summing up the spiking activities of MNs, forming the cumulative spike train (CST). The CST reflects the neural drive transmitted to the innervated muscle (Farina and Negro, 2015). Interestingly, the CST, and so the common synaptic input, carries oscillations at least up to 75 Hz (Muceli et al., 2022), but only frequencies below 10 Hz are transduced effectively into muscle force changes. Oscillations with frequencies at about 10 Hz and higher are filtered out by spinal networks (Williams and Baker, 2009; Williams et al., 2010) and the low-pass filtering characteristics of the muscle (Mannard and Stein, 1973; Baldissera et al., 1998; Farina et al., 2014). Oscillatory activities found in the beta (13 Hz to 30 Hz) and gamma band (30 Hz to 70 Hz) during tonic and dynamic muscle contractions, respectively, are at least partly cortical in origin, as shown by connectivity analyses like cortico-muscular coherence (CMC) (Conway et al., 1995; Halliday et al., 1998; Farmer, 1998; Marsden et al., 2000; Omlor et al., 2007; Gwin and Ferris, 2012; Ibáñez et al., 2021). The functional interpretation of these high-frequency oscillations is unclear as they do not directly affect muscle force. They may serve as a probing signal to the periphery, facilitate a steady motor output, or relate to motor readiness (Baker, 2007; Engel and Fries, 2010; Jenkinson and Brown, 2011). An additional hypothesis, which is not mutually exclusive, is that some CST components in the beta or gamma band represent oscillations in cortical areas that propagate downstream to MNs – potentially via corticomotoneuronal connections (Brouwer and Ashby, 1992; Maertens De Noordhout et al., 1999; Lemon, 2008; Ibáñez et al., 2021) – without serving a specific function in those neurons. We refer to this type of propagation as *leaking*. Since beta and gamma bands are not within the effective musculoskeletal bandwidth, the spinal networks do not have to implement any filtering or cancellation strategies to prevent interference with muscle control signals. If this leaking-oscillations hypothesis is true, the *modulations* of such leaking cortical oscillations should be visible in the CST of the downstream MNs. To support or contest this hypothesis, we investigated in this study whether the power modulation of cortical oscillations *unrelated* to an executed movement is paralleled by the same modulations in the CST of spinal MNs. We experimentally manipulated cortical oscillations of human participants indirectly by asking them to perform various mental tasks (foot motor imagery, hand motor imagery, mental arithmetic, and no task) while simultaneously executing a static isometric dorsiflexion of the right foot (see Figure 1a). The employed mental tasks are known to induce specific macroscale cortical oscillation patterns, which are measurable in electroencephalography (EEG) signals (Pfurtscheller and Lopes da Silva, 1999; Neuper et al., 2005; Friedrich et al., 2012). Importantly, the imagined movements were different from the executed isometric dorsiflexion of the right foot that was necessary to generate MN activity.

**Figure 1.**
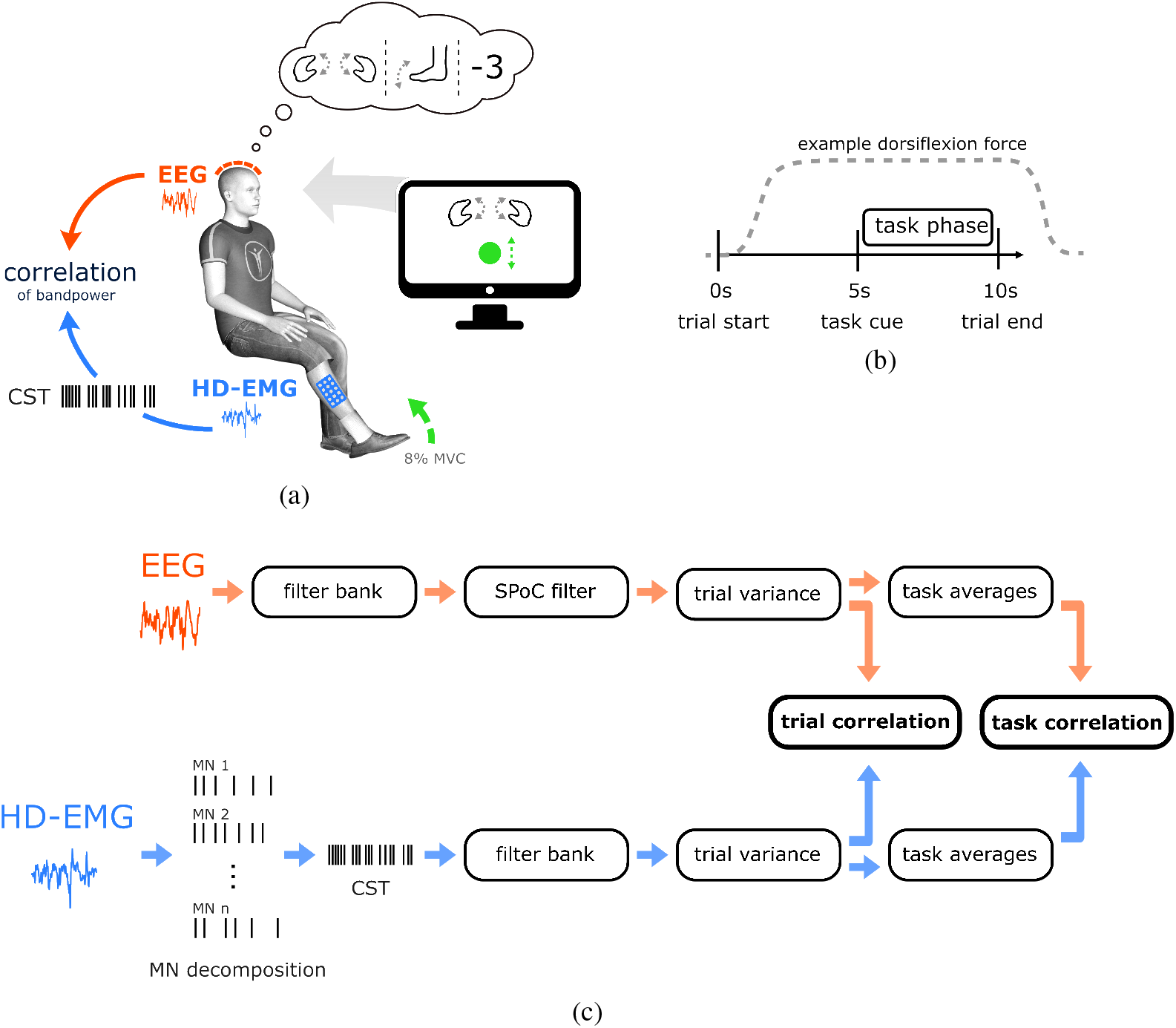
**a:** Experiment setup. HD-EMG and EEG were recorded while participants performed an isometric dorsiflexion at 8% MVC together with a mental task (hand MI, foot MI, mental arithmetic, or no task). The computer screen displayed task instructions and provided feedback on the dorsiflexion force. **b:** Sequence of a trial. Participants increased the dorsiflexion force to 8% MVC at the beginning of a trial, and the mental tasks were performed during the task phase. **c:** Overview of the *trial* and *task* correlation analyses. The cummulative spike train (CST) of spinal motor neurons was extracted from HD-EMG signals. The EEG and CST trial variances were calculated over the task phase, yielding band-power values which were then correlated across individual trials and task averages.

We hypothesized that power modulations of cortical oscillations induced by these mental tasks leak downstream to MNs, and are detectable in their CST. The CST activity was obtained by decomposing the MN activity from high-density electromyography (HD-EMG) signals recorded non-invasively from the right tibialis anterior (TA) muscle (Negro et al., 2016). We calculated CST and EEG power across frequency bands within trials, and analyzed the correlations between CST and EEG band power across *trials* and *mental tasks* to identify leaking oscillations (see Figure 1a and 1c).

Cortical oscillations induced by mental tasks that leak downstream and do not interfere with the motor output could allow the implementation of a control signal for neural interface applications, analogous to oscillation-based brain-computer interfaces (BCIs) (Friedrich et al., 2013). A control signal derived from leaking oscillations could also allow the extension of the degrees-of-freedom (DoF) of the human body without affecting existing movement functionality (Eden et al., 2022), allowing movement augmentation scenarios. To assess the potential of such a control approach, we evaluated the discriminability of the mental tasks from the band power of CST oscillations using signal-to-noise ratio (SNR) analysis and single-trial classification.

## 2 Results

### 2.1 Power Comodulation

We calculated the Kendall rank correlation coefficient *τ* between the band powers of CST and EEG to reveal a potential power comodulation between MN and brain activity. Band power was calculated from 5 Hz wide bands, separated by 1 Hz steps (i.e., 5-9, 6-10, …, 70-74 Hz). EEG signals were first mapped to a one-dimensional space using a data-driven spatial filter (Source Power Comodulation (SPoC) (Dähne et al., 2014)) before EEG band power was calculated.

The correlation between CST and EEG band power was computed (1) across all *trials* irrespective of the task as *trial correlation* (*τ*_*trials*_), and (2) across *tasks* using the median power values of each task as *task correlation* (*τ*_*tasks*_). Figure 2a shows the group average of the trial and task correlations and their 95% confidence intervals as a function of frequency. The task correlations were noisy due to the small number of data points per participant (4 data points or tasks). For visualization purposes, we therefore smoothed the task correlations before averaging over participants with a moving average comprising 10 frequency bands. The group average of the trial correlations peaked in the beta band at 24 Hz with a value of 0.08. The group average of the smoothed task correlations showed a peak at 31 Hz with a value of 0.14. Both trial and task correlations displayed increased correlations in the beta range from approximately 15 to 25 Hz. Moreover, the task correlation was increased in the lower gamma band around 30 Hz. The positive *task correlations* indicate that the mental tasks modulated the EEG and CST band power signals in the same direction. In addition, the prominent positive *trial correlations* suggest that there were *spontaneous* trial-by-trial correlations between EEG and CST band power signals that were not caused by the mental tasks.

**Figure 2.**
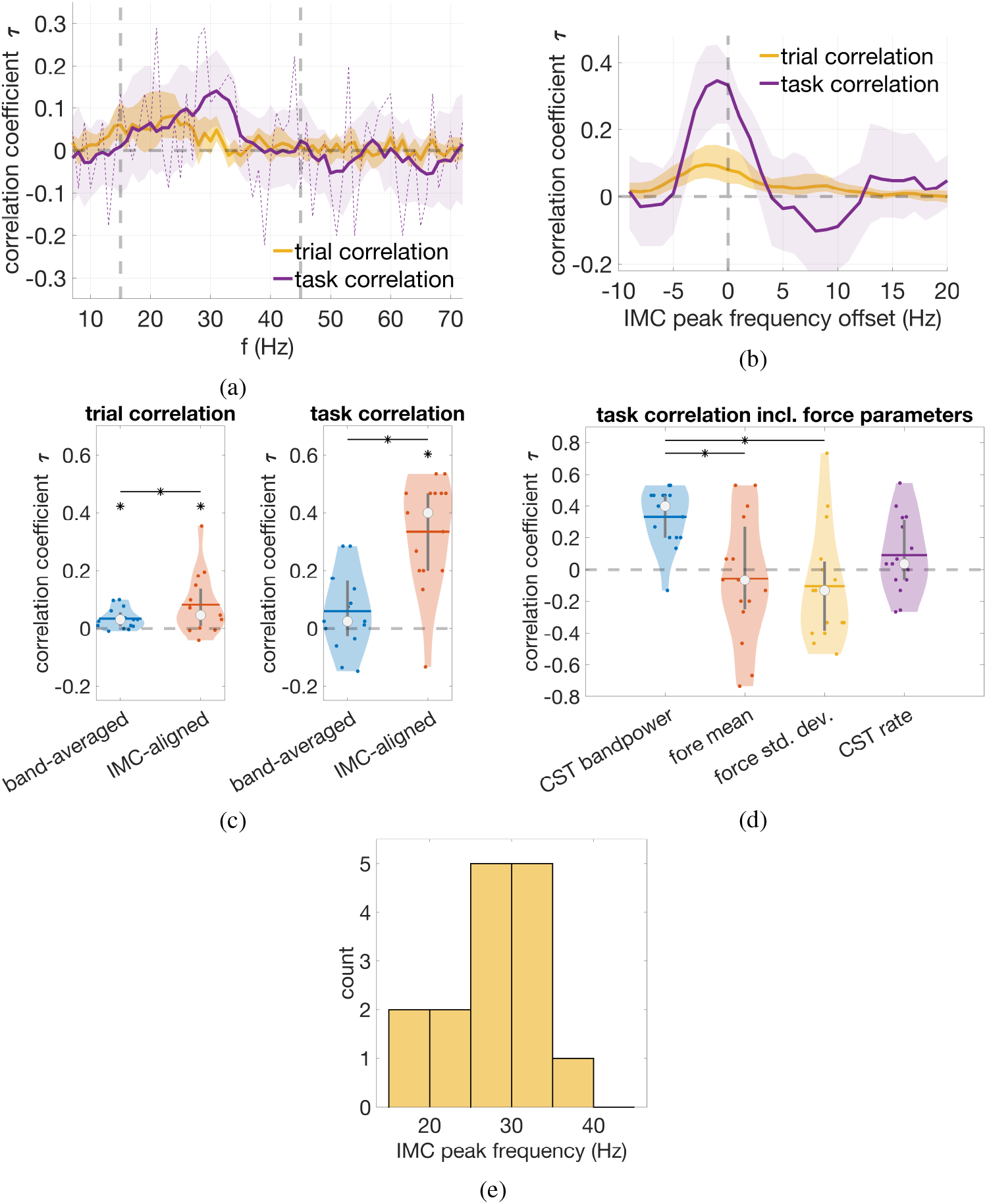
CST-EEG correlation results. **a:** Trial and task correlations as a function of frequency. Shown are group averages and their 95% confidence intervals. The task correlations were smoothed by a moving average filter across 10 frequency bands; the unsmoothed task correlations are shown by a purple dashed line. The vertical dashed lines mark the frequency range used for calculating the band-averaged correlations. **b:** IMC-aligned trial and task correlations over IMC peak frequency offsets. Shown are group averages and their 95% confidence intervals. **c:** Band-averaged and IMC-aligned trial and task correlations of the individual participants. The group medians are marked with a circle; the group mean values are marked with a horizontal line; the vertical grey lines extend from the 1^st^ to the 3^rd^ quartile. A star marks statistically significant differences between groups or against 0 (two-sided Wilcoxon signed-rank test, *q* = 0.05). **d:** Comparison of IMC-aligned task correlations based on CST bandpower features and force-related parameters (force mean, force std. dev., CST rate). All task correlations were computed against SPoC-filtered EEG. A star marks statistically significant differences between groups (Tukey-Kramer post-hoc test, *α* = 0.05). **e:** Distribution of the found IMC peak frequencies.

The correlations were then averaged over the beta and low-gamma bands (15 Hz to 45 Hz), which we refer to as *band-averaged* correlation. In addition, we determined the participant-specific frequency with the largest common synaptic input, as measured by intramuscular coherence (IMC), and averaged the correlations around the IMC peak (*IMC-aligned* correlation). The *band-averaged* and *IMC-aligned* correlations are shown in Figure 2c and in Table 1. A two-sided Wilcoxon signed-rank test was used to test for significant differences, and p-values were adjusted for multiple comparisons. Both band-averaged and IMC-aligned *trial* correlations were significantly greater than 0 (*W* = 113, *p*_*adj*_ = 0.002, and *W* = 109, *p*_*adj*_ = 0.005, respectively). Furthermore, the IMC-aligned trial correlation was significantly larger than the band-averaged correlation (*W* = 23, *p*_*adj*_ = 0.042). The IMC-aligned *task* correlation was significantly larger than both 0 and the band-averaged correlation (*W* = 118.5, *p*_*adj*_ = 0.001, and *W* = 0, *p*_*adj*_ = 4e*−*4, respectively). In contrast, the band-averaged task correlation was not significantly larger than 0 (*W* = 88.5, *p*_*adj*_ = 0.110). Thus, the IMC alignment increased the trial and task correlations relative to the broadband average.

**Table 1:**
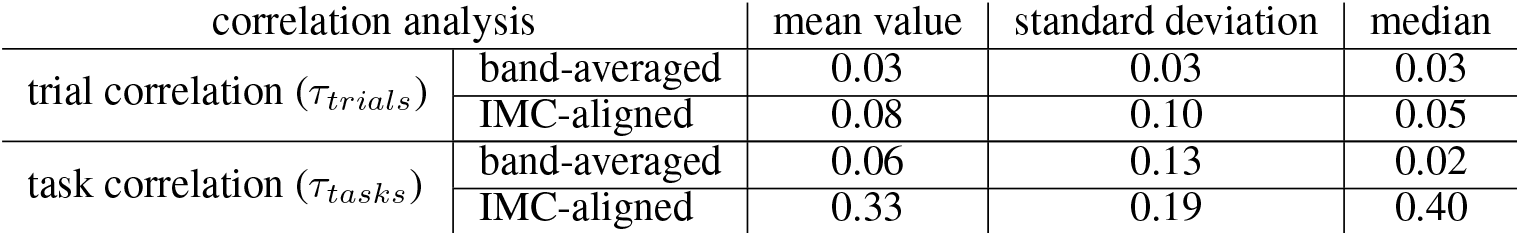
Group statistics of the Kendall rank correlation coefficient τ across trials and tasks.

The event-related desynchronization/synchronization (ERD/S) analysis in Section 2.5 revealed that only the active mental tasks, i.e., *foot MI, hand MI*, and *math*, featured clear relative power decreases in the mu and beta bands, while the *no task* condition did not. The reported power comodulations could be solely due to bandpower differences between *no task* and *any* other task, rather than bandpower differences within the other tasks. We therefore analyzed whether there were significant power comodulations within the task subset comprising *foot MI, hand MI*, and *math*. For this purpose, we used the same trained spatial SPoC filters but evaluated the power correlations using only the task subset. The obtained group-level correlations are shown in Table 2. The IMC-aligned task correlation was significantly larger than both 0 and the band-averaged correlation (*W* = 116, *p*_*adj*_ = 0.002, and *W* = 6, *p*_*adj*_ = 0.002, respectively). The band-averaged correlation was also significantly larger than 0 (*W* = 98.5, *p*_*adj*_ = 0.032). All tests were two-sided Wilcoxon signed-rank tests, and p-values were adjusted for multiple comparisons. This indicates that the observed power comodulations were not solely due to greater bandpower in *no task* compared to the other tasks, and that *foot MI, hand MI*, and *math* yielded common power changes in EEG and CST.

**Table 2:** Group statistics of the Kendall rank correlation coefficient *τ* across the task subset. Only the mental tasks *foot MI, hand MI*, and *math* were considered.

| correlation analysis |  | mean value | standard deviation | median |
| --- | --- | --- | --- | --- |
| task correlation ($\tau_{tasks}$) | band-averaged | 0.09 | 0.14 | 0.06 |
|  | IMC-aligned | 0.39 | 0.28 | 0.47 |

To assess whether task-dependent force changes could have confounded the results, we calculated task correlations using different force-related parameters instead of CST bandpower, namely force mean, force std. dev., and CST rate. We trained the SPoC filters using the individual force-related parameters as target signals, analogously to the procedure described in Section 5.6 but without removing force-related confounders. The task correlations at the IMC peak (IMC aligned) are shown in Figure 2d, along with the task correlation for CST bandpower reported previously. In terms of mean and median values, the task correlation of CST bandpower was the highest, and the task correlations of the force-related parameters were close to zero. We compared task correlations using a one-way repeated measures ANOVA with target signal type as the explanatory variable, and found a statistically significant difference between task correlations (*F* (3, 42) = 6.868, *p* = 0.001). A Tukey-Kramer post-hoc test revealed statistically significant pairwise differences between CST bandpower and force mean (*p* = 0.044, 95%-CI = [0.0%, 0.8%]), and CST bandpower and force std. (*p* = 0.002, 95%-CI = [0.2%, 0.7%]). CST bandpower and CST rate were not statistically significantly different (*p* = 0.064, 95%-CI = [0.0%, 0.5%]). Overall, these results support the interpretation that the observed CST-EEG bandpower correlations were rather due to cortical oscillations leaking downstream than a spurious correlation due to task-dependent force output as a confounding factor.

The IMC peak was used to determine the frequency with the largest common synaptic input (Farina et al., 2014). Figure 2e displays the histogram of the found peak frequencies. The majority of peaks were found in the upper-beta/lower-gamma band around 30 Hz, which is also the frequency range yielding the highest group task correlation. However, participants P5, P7, and P11, which exhibited greater task correlation, had IMC peak frequencies in the lower-beta band around 20 Hz (c.f. Figure 3). Figure 2b shows trial and task correlations as a function of an offset to the IMC peak frequency. The highest average trial and task correlations were around an offset of *−*2 Hz and *−*1 Hz, respectively.

**Figure 3.**
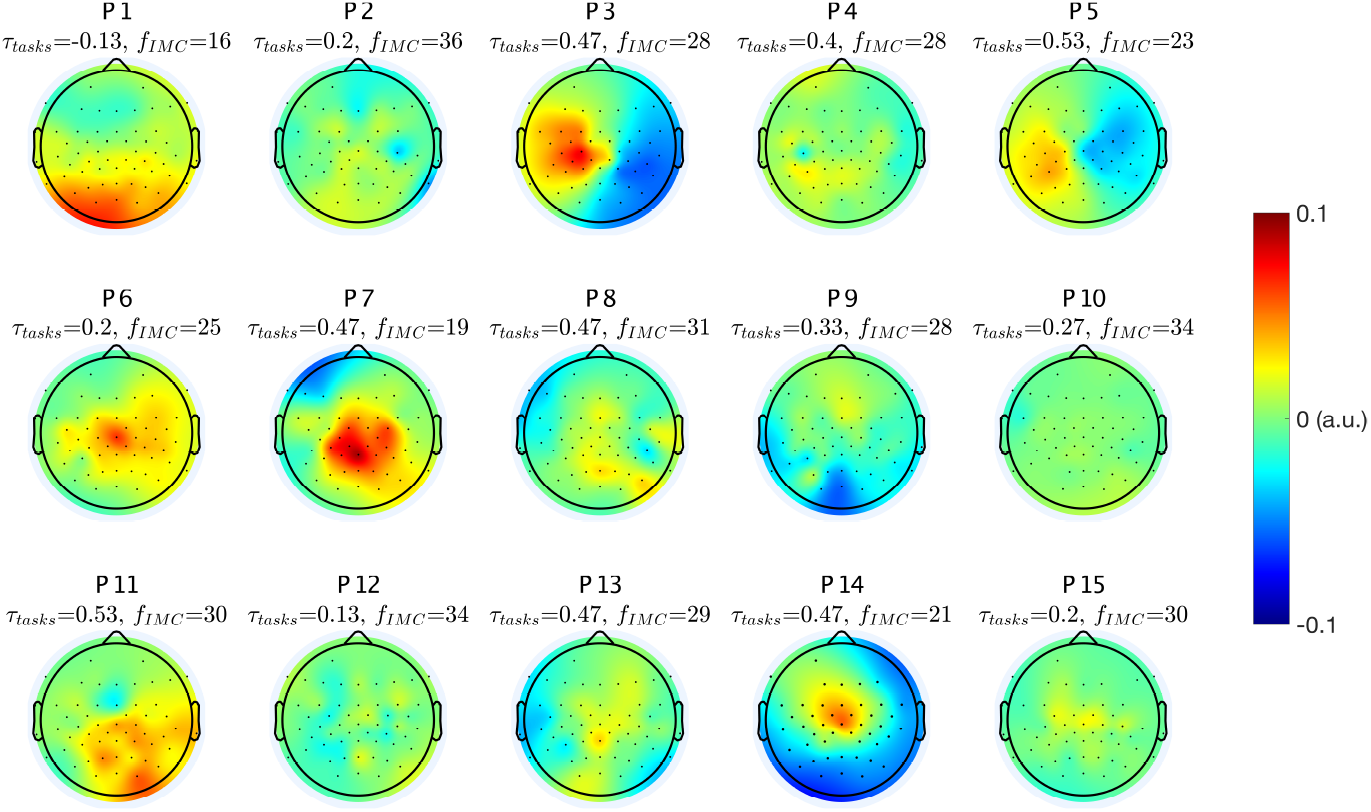
SPoC pattern of each participant and the IMC peak frequency *f*_*IMC*_ in Hz with the respective task correlation coefficient *τ*_*tasks*_ at this frequency.

Thus, the highest correlations were obtained around the IMC peak frequency, suggesting that the CST-EEG band power correlations are primarily due to oscillations carried by the common synaptic input.

SPoC finds a spatial filter that extracts a signal whose power has maximal covariance with a target variable (Dähne et al., 2014). One can find the corresponding *pattern* to this spatial filter. The pattern displays the source generating that signal. Figure 3 shows the channel-space SPoC patterns at the participant-specific IMC peaks and the associated task correlations *τ*_*tasks*_ and IMC peak frequencies. Participants P3, P5, P6, P7, and P14 displayed a pattern that suggests a dipole-like source located close to the foot area of the primary motor or somatosensory cortex. P1 and P11 displayed an occipital/parietal source, with only P11 having a greater correlation. Notably, P3, P5, P7 and P14 achieved relatively high correlation and classification accuracies (c.f. Table 4). Not all participants exhibited distinct and clear SPoC patterns. This is possibly due to the noisy nature of the EEG signal and the imperfect transmission of oscillations downstream to MNs, which affects the target signal used to train the SPoC model and thus the model itself.

### 2.2 Signal-to-Noise Ratio Analysis

An SNR analysis was performed to analyse the discriminability of CST band power with regard to mental tasks. The group-level SNR of CST band power as a function of frequency is shown in Figure 4a. As SNR values are not limited, a robust group-level average was calculated using the median. For visualization purposes, we smoothed the SNR values before computing the group median with a moving average comprising 5 frequency bands. Note that due to the bias removal and statistical fluctuations, negative SNR values can occur.

**Figure 4.**
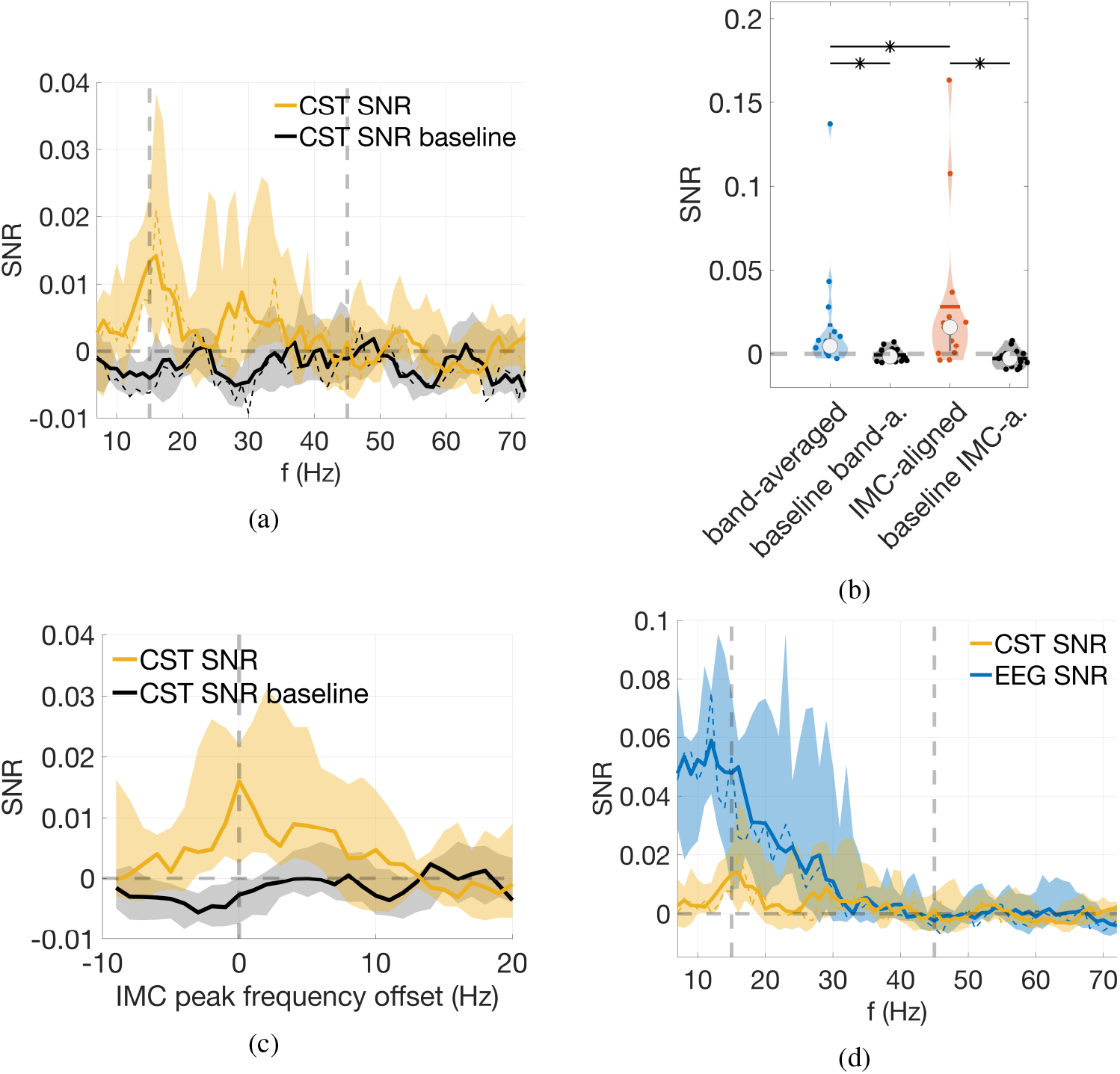
SNR results. **a, c**, and **d** show group medians and their 95% confidence intervals. **a:** CST and baseline SNR as a function of frequency. SNR values were smoothed with a moving average filter using 5 frequency bands; the unsmoothed SNR values are shown as dashed lines. The vertical dashed lines mark the frequency range used to calculate the band-averaged SNR. **b:** Band-averaged and IMC-aligned CST SNR of the individual participants with their corresponding baselines. The group medians are marked with a circle; the mean values are marked with a horizontal line; the vertical grey lines extend from the 1^st^ to the 3^rd^ quartile. A star marks statistically significant differences (two-sided Wilcoxon signed-rank test, *q* = 0.05). **c:** IMC-aligned CST SNRs as a function of an offset to the IMC peak frequency. **d:** CST SNR and EEG SNR. The latter was derived from electrode Cz.

One can visually identify CST SNR peaks around 15 and 30 Hz, i.e., in beta and low-gamma bands. We show band-averaged (15 Hz to 45 Hz) and IMC-aligned CST SNRs in Figure 4b and Table 3. The band-averaged and IMC-aligned CST SNRs were both statistically significantly larger than their baselines (*W* = 111, *p*_*adj*_ = 0.003, and *W* = 114, *p*_*adj*_ = 0.003, for band-averaged and IMC-aligned CST SNR, respectively). Moreover, the IMC-aligned CST SNR was statistically significantly larger than the band-averaged CST SNR (*W* = 12, *p*_*adj*_ = 0.004). All tests were two-sided Wilcoxon signed-rank tests, and p-values were adjusted for multiple comparisons. The effect of a frequency offset on the IMC-aligned CST SNR is shown in Figure 4c. The CST SNR peaked around the original IMC peak frequency (offset of 0 Hz), but this peak was not clearly distinct.

**Table 3:** Group-level CST SNR.

|  | mean (%) | std. dev. (%) | median (%) |
| --- | --- | --- | --- |
| band-averaged | 0.02 | 0.04 | 0.00 |
| baseline band-averaged | 0.00 | 0.00 | 0.00 |
| IMC-aligned | 0.03 | 0.05 | 0.02 |
| baseline IMC-aligned | 0.00 | 0.01 | 0.00 |

**Table 4:** Classification accuracies in % for each participant obtained with different features.

|  | P1 | P2 | P3 | P4 | P5 | P6 | P7 | P8 | P9 | P10 | P11 | P12 | P13 | P14 | P15 |
| --- | --- | --- | --- | --- | --- | --- | --- | --- | --- | --- | --- | --- | --- | --- | --- |
| CST-FB | 31 | 28 | 32 | 32 | 43 | 24 | 38 | 26 | 29 | 27 | 28 | 29 | 27 | 30 | 30 |
| CST-IMC | 30 | 26 | 32 | 31 | 43 | 24 | 37 | 26 | 29 | 27 | 29 | 29 | 27 | 30 | 31 |
| FORCE-MEAN | 34 | 31 | 30 | 31 | 32 | 26 | 28 | 25 | 27 | 36 | 24 | 32 | 28 | 28 | 24 |
| FORCE-STD | 27 | 25 | 30 | 27 | 30 | 27 | 31 | 30 | 26 | 24 | 24 | 24 | 27 | 27 | 22 |
| CST-RATE | 27 | 27 | 27 | 29 | 30 | 29 | 31 | 25 | 25 | 23 | 25 | 22 | 28 | 27 | 20 |
| EEG-FB | 42 | 38 | 38 | 38 | 52 | 35 | 41 | 34 | 37 | 30 | 29 | 34 | 31 | 41 | 29 |

Finally, we compared CST and EEG SNR profiles in Figure 4d. Even though only data from the Laplace filtered EEG channel Cz was used, the EEG SNR was prominent and higher than the CST SNR. This confirms that the employed mental tasks can be detected in macroscale brain oscillations over the motor cortex while an isometric movement is being performed. The EEG SNR peaked in the mu band around 12 Hz and yielded pronounced values up to around 30 Hz. Thus, the motor cortex carries task informative oscillations, which could leak down to MNs. We chose to use a Laplace filter instead of SPoC because SPoC is a supervised method and depends on the discriminability of the target signal, i.e, the CST. It would result in an ineffective spatial EEG filter at frequencies with a low CST SNR.

### 2.3 Classification

We extracted features from CST, EEG and potential confounders from the task phase of trials. The CST and EEG features, *CST-FB* and *EEG-FB*, were bandpower features computed using a filter bank (15-19, 17-21, …, 41-45 Hz). In addition, we computed CST bandpower features that were aligned with the participants’ IMC peaks (*CST-IMC*). Both *CST-FB* and *CST-IMC* features potentially capture cortical oscillations that leak downstream. The potential confounder features were computed as the mean and standard deviation of the dorsiflexion force and the CST rate (*FORCE-MEAN, FORCE-STD* and *CST-RATE*). The four mental tasks were then classified with a shrinkage linear discriminant analysis (sLDA) classifier (Peck and Van Ness, 1982; Blankertz et al., 2011) using 10 × 10 fold cross-validation. The classification accuracies and group statistics are provided in Figure 5a and Tables 4 and 5. The classification accuracies yielded by CST-FB, CST-IMC, FORCE-MEAN, FORCE-STD and EEG-FB were statistically significantly higher than the chance level of 25% (*W* = 118, *p*_*adj*_ = 4e−4; *W* = 119, *p*_*adj*_ = 4e−4; *W* = 112, *p*_*adj*_ = 0.002; *W* = 96, *p*_*adj*_ = 0.050; *W* = 120, *p*_*adj*_ = 4e−4). The classification accuracy of CST-rate was not statistically significant (*W* = 89, *p*_*adj*_ = 0.107). All tests were two-sided Wilcoxon signed-rank tests, and p-values were adjusted for multiple comparisons. The statistical tests show that the oscillations in the CST and EEG signals were modulated by the mental tasks. No strong preference for any particular mental task was present in the confusion matrix for the CST-FB features (Figure 5c).

**Table 5:** Group statistic of the classification accuracies obtained with different features.

|  | mean (%) | std. dev. (%) | median (%) |
| --- | --- | --- | --- |
| CST-FB | 30 | 5 | 29 |
| CST-IMC | 30 | 5 | 29 |
| FORCE-MEAN | 29 | 4 | 28 |
| FORCE-STD | 27 | 3 | 27 |
| CST-RATE | 26 | 3 | 27 |
| EEG-FB | 37 | 6 | 37 |

**Figure 5.**
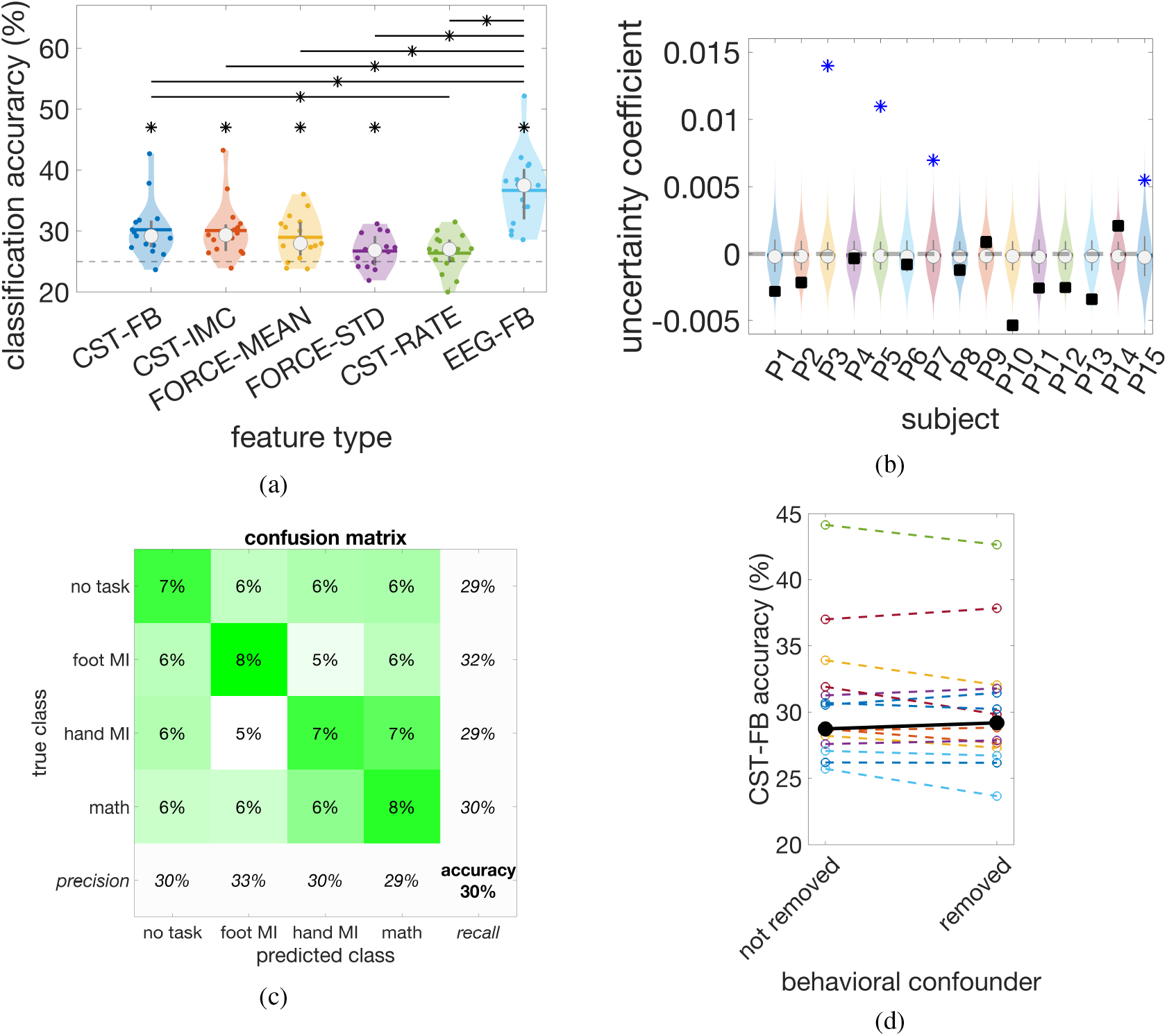
Classification results. **a:** Classification accuracies obtained from different features. Each dot represents a participant. The horizontal dashed line indicates the chance level of 25%. The group medians are marked with a circle; the group mean values are marked with a horizontal line; the vertical grey lines extend from the 1^st^ to the 3^rd^ quartile. A star marks statistically significant differences from the chance level or between groups (two-sided Wilcoxon signed-rank test with *q* = 0.05 or Tukey-Kramer post-hoc test with *α* = 0.05, respectively). **b:** Uncertainty coefficients *U* (*X, Y*) between the class predictions from CST-FB and EEG-FB features are shown as black boxes; statistically significant values are indicated with a blue asterisk (one-sided permutation test, *q* = 0.05). Additionally, we show the distribution of the uncertainty coefficients when permuting class labels as colored violin plots. **c:** Confusion matrix including recall, precision and accuracy values on the sides. The coloured true/predicted fields sum up to 100%. Trials from all participants were pooled. **d:** Comparison of CST-FB classification accuracies before and after confounder removal with regression (i.e., classification of residuals). The black dots indicate the medians.

The mental tasks also induced task-dependent changes in force and CST rate features. These task-dependent changes could potentially have confounded the classification analysis of the CST band-power features. Figure 5d shows the CST-FB classification accuracies with and without the removal of these potential confounders using a multiple linear regression (see Methods for details). The removal of these potential confounders did not have a statistically significant effect on the group-level classification accuracy (*W* = 85, *p* = 0.169, two-sided Wilcoxon signed-rank test). Furthermore, we calculated the median across the pairwise classification accuracy differences (i.e., confounders *not removed* minus *removed*). This yielded a median difference of 0.4% with a 95% confidence interval of CI = [−0.5%, 1.1%]. The narrow confidence interval around 0% indicates that the removal of potential confounders had a minor impact on the classification accuracies. Thus, it is unlikely that force changes confounded the classification accuracy obtained from CST band-power features.

The classification accuracies were then compared using a one-way repeated measures ANOVA with feature type as the explanatory variable. A statistically significant difference between classification accuracies was found (*F* (5, 70) = 21.452, *p* = 2e*−*8, GG adjusted). A Tukey-Kramer post-hoc test revealed statistically significant pairwise differences between CST-FB and CST-RATE (*p* = 0.048, 95%-CI = [0.0%, 7.6%]), EEG-FB and CST-FB (*p* = 2e*−*4, 95%-CI = [3.2%, 9.7%]), EEG-FB and CST-IMC (*p* = 3e*−*4, 95%-CI = [3.1%, 10.1%]), EEG-FB and FORCE-MEAN (*p* = 0.002, 95%-CI = [2.8%, 12.6%]), EEG-FB and FORCE-STD (*p* = 2e*−*5, 95%-CI = [5.7%, 14.1%]), and EEG-FB and CST-RATE (*p* = 1e*−*5, 95%-CI = [6.2%, 14.4%]).

Mental tasks could be classified from force-related features (force mean value, force standard deviation, and CST rate). We investigated whether there was a correlation between the classification accuracies yielded by CST band-power features and force-related features (Figure 6a-6c). Importantly, the effect of force and CST rate on the CST-FB features was not removed here to reveal any potential correlation of the raw features. We provide the Kendall rank correlation coefficients *τ* of the classification accuracies, associated p-values (two-sided permutation test), and 95% confidence intervals in the plots. The results did not indicate any apparent correlation of classification accuracies between CST-FB features and the three force-related features. In line with that, the Kendall rank correlation coefficients were not statistically significant. Note, however, that confidence intervals were large, and, thus, the available data could not demonstrate the absence of correlation. Between CST and EEG band-power features, a statistically significant correlation of *τ* = 0.45 (*p* = 0.021) was observed (Figure 6d CST-FB vs. EEG-FB).

**Figure 6.**
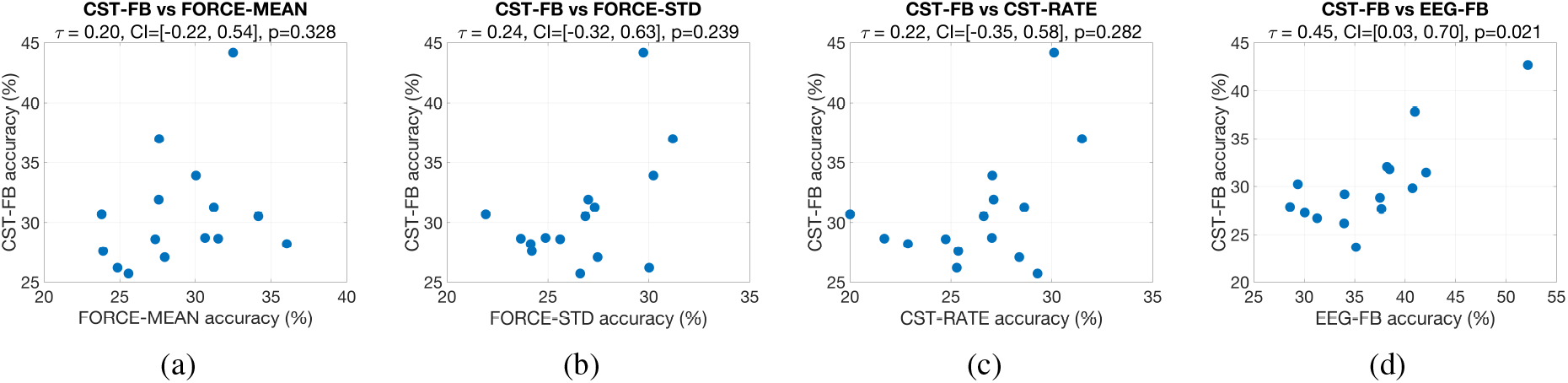
Correlation plot of classification accuracies. The Kendall rank correlation coefficient *τ* of the classification accuracies and its 95% confidence interval and p-value are shown in the title. **a-c:** Comparisons of features capturing force output **d:** Comparison with EEG (note the altered scale).

We further analysed the association between the predicted class labels from CST-FB and EEG-FB features with the uncertainty coefficient *U* (*X, Y*) (Press et al., 2007) (see Figure 5b). Statistically significant coefficients were obtained for participants P3 (*p*_*adj*_ = 1e*−*4), P5 (*p*_*adj*_ = 1e*−*4), P7 (*p*_*adj*_ = 1e*−*4), and P15 (*p*_*adj*_ = 0.034) (one-sided permutation tests adjusted for multiple comparisons across subjects at *q* = 0.05). Thus, for these participants, the brain signals and the CST yielded related class predictions for individual trials.

### 2.4 Analysis of Force-related Parameters

We analysed four parameters reflecting force changes, i.e., force mean, force std. dev., CST rate and bipolar EMG variance. The bipolar EMG variance was derived from an antagonist muscle (medial head of the right gastrocnemius muscle) after 10 Hz high-pass filtering. The parameter values are shown in Figure 7 for the investigated mental tasks. Figure 7a suggests that participants applied a marginally greater force during the math task compared to the other tasks. The group average for *math* was up to 0.05% higher than for the other tasks. Interestingly, the force in the other tasks *no task, foot MI*, and *hand MI* was, on average, between 0.03% and 0.04% below the target force. In contrast, the group average in the math task was 0.01% above the target force. No apparent differences were observed for the other parameters (Figure 7b to 7d).

**Figure 7.**
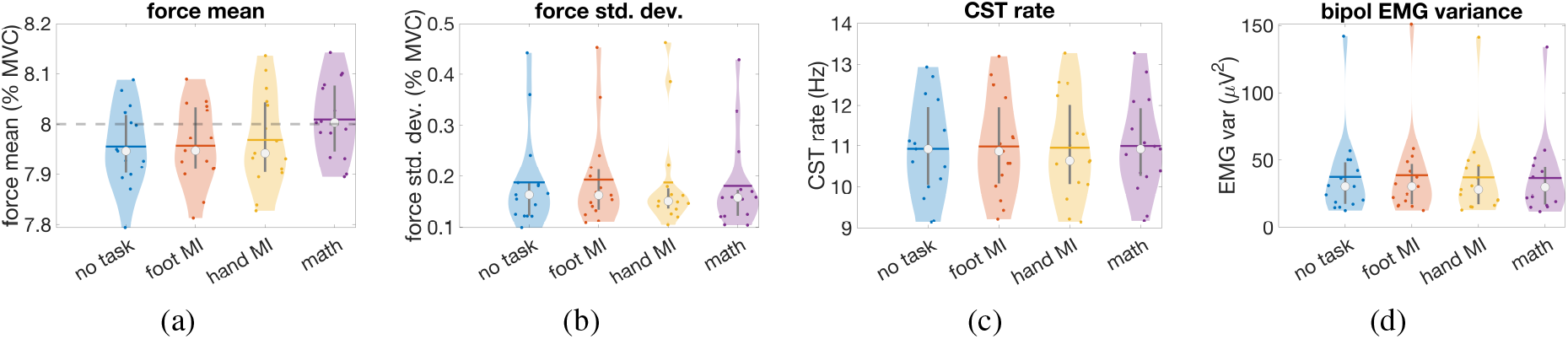
Force-related parameters during the task phase (6 s to 9.5 s after trial start) as a function of the mental task. The data points are the participants’ medians over trials for the respective parameter and task. The group medians are marked with a circle; the group mean values are marked with a horizontal line; the vertical grey lines extend from the 1^st^ to the 3^rd^ quartile. **a:** Mean value of the force. The horizontal dashed line marks the target force at 8%. **b:** Standard deviation of the force. **c:** CST rate. **d:** EMG variance of an antagonist muscle (medial head of the right gastrocnemius muscle).

The standard deviation of the force was high for three participants in each task (see Figure 7b). For *no task, hand MI*, and *math*, these participants were consistently P6, P15, and P11, listed in order of decreasing standard deviation. In the case of *foot MI*, P6, P15 and P7 had the highest standard deviation. P6, P15, and P11 did not consistently achieve a high classification accuracy for CST-FB or FORCE-STD features (c.f. Table 4). Only P7 achieved a notably high classification accuracy for CST-FB at 38% (31% for FORCE-STD).

We compared the force-related parameters shown in Figure 7 across the mental tasks. Each parameter was treated as a separate response variable, and we conducted four separate one-way repeated measures ANOVAs with mental task as the explanatory variable. The p-values were adjusted for multiple comparisons. A statistically significant difference between mental tasks was found for force mean (*F* (3, 42) = 12.379, *p*_*adj*_ = 2e−5), whereas no significant differences were found for force std. dev. (*F* (3, 42) = 1.565, *p*_*adj*_ = 0.283), CST rate (*F* (3, 42) = 0.959, *p*_*adj*_ = 0.421), or bipolar EMG variance (*F* (3, 42) = 2.882, *p*_*adj*_ = 0.182, GG adjusted).

### 2.5 EEG ERD/S Plots

The event-related desynchronization/synchronization (ERD/S) plots for the four mental tasks are shown in Figure 8. We computed the ERD/S using an STFT with a 1-second Hanning window and the gain model (Grandchamp and Delorme, 2011; Delorme and Makeig, 2004). The plots depict the median across all participants of the Laplace-filtered central EEG electrode Cz. The baseline interval ranged from 3 s to 5 s and included the isometric dorsiflexion of the right foot. This minimized the impact of the executed isometric movement on the ERD/S plots.

**Figure 8.**
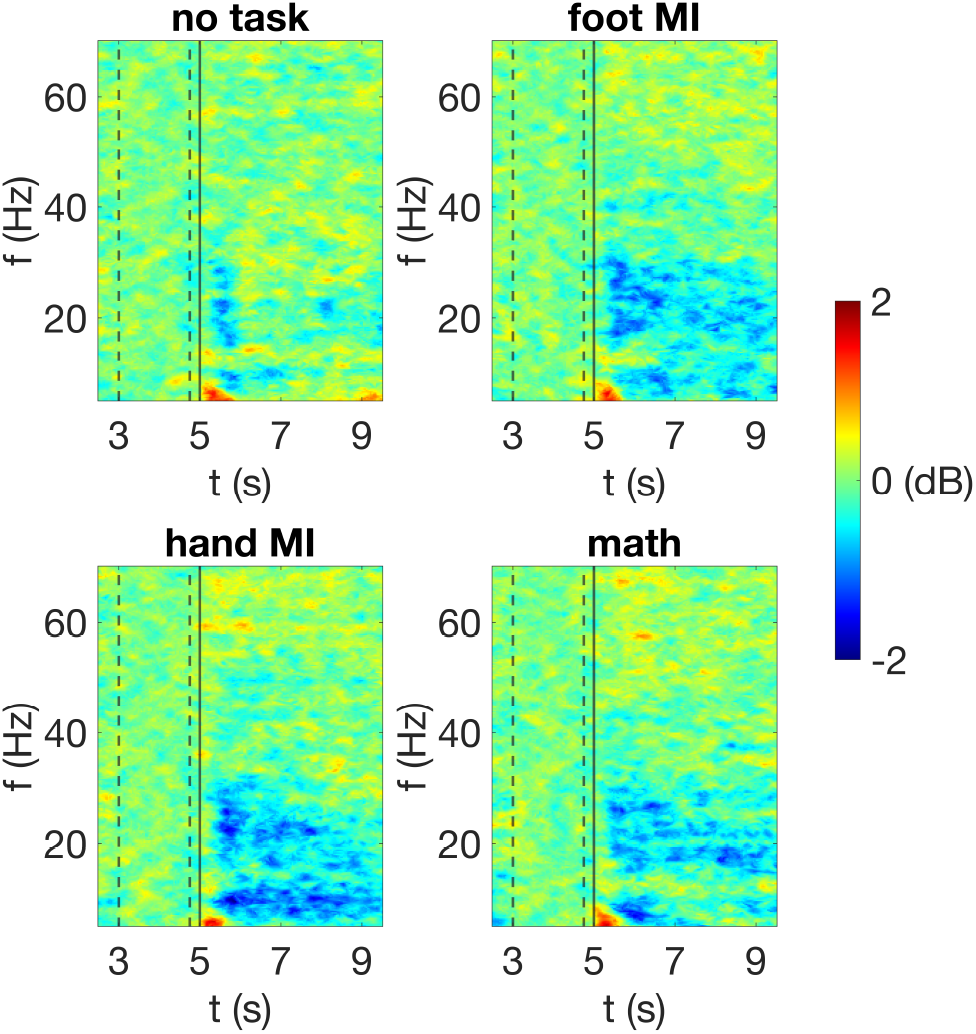
Group-level ERD/S maps of the four mental tasks at electrode Cz. The task cue was at second 5. Colors depict power relative to the baseline interval ranging from 3 s to 5 s as indicated by the dashed lines.

A relative power decrease, i.e., ERD, occurred shortly after cue presentation in the mu and beta bands for all four tasks. In the *no task* condition, there was no pronounced ERD or ERS observable after 6 s. In contrast, *foot MI, hand MI*, and *math* exhibited an ERD in the beta band between 15 Hz to 30 Hz. Additionally, there was an ERD in the mu band around 10 Hz for *foot MI* and particularly *hand MI*. The ERD shortly after cue presentation could reflect the processing of the external stimulus. The pronounced ERD in *foot MI, hand MI*, and *math* after 6 s is likely related to the execution of the respective mental task.

## 3 Discussion

We found that *power modulations* of cortical beta and low-gamma oscillations induced by mental tasks were accompanied by concordant power modulations in the cumulative spike train of spinal motor neurons. The mental tasks were unrelated to the executed movement necessary to create CST activity (sustained dorsiflexion). Furthermore, the observed correlations were not explained by potential force-related confounders, and the identified EEG sources mostly aligned with the foot area on the primary motor or somatosensory cortex. These findings support the hypothesis that *force-unrelated* cortical oscillations above 10 Hz can leak downstream to MNs and are not removed by spinal networks.

### Connectivity Analyses

The transmission of beta and low-gamma oscillations from the motor cortex to MNs is well-documented with connectivity analyses such as corticomuscular coherence (Conway et al., 1995; Halliday et al., 1998; Farmer, 1998; Marsden et al., 2000; Omlor et al., 2007; Gwin and Ferris, 2012; Ibáñez et al., 2021), directed coherence (Witham et al., 2011) or directed transfer function (Tsujimoto et al., 2009). Furthermore, Bräcklein et al. discovered that beta bursts at the cortical and MN levels have the same source, suggesting that MN beta oscillations result from cortical projections (Bräcklein et al., 2022), such as corticomotoneuronal connections (Brouwer and Ashby, 1992; Maertens De Noordhout et al., 1999; Lemon, 2008). It also has been found that the simultaneous execution of additional motor or non-motor tasks reduces CMC (Kristeva-Feige et al., 2002; Mat Safri et al., 2007; Johnson et al., 2011). However, in contrast to previous studies, we were not investigating EEG-CST connectivity per se, but whether brain and MN oscillations comodulate in power across mental tasks due to the existing connectivity. In contrast to our findings, (Kristeva-Feige et al., 2002; Mat Safri et al., 2007; Johnson et al., 2011) did not observe spontaneous or task-related band-power comodulations between EMG and EEG signals. This could be because EMG signals suffer from interferences between motor unit action potentials, potentially masking bandpower changes, while the CST does not. Moreover, we employed a supervised data-driven spatial EEG filter, which typically offers better SNR compared to a fixed spatial filter. Furthermore, while (Kristeva-Feige et al., 2002; Mat Safri et al., 2007; Johnson et al., 2011) recorded EMG from hand muscles, we derived MN activities from the TA muscle. It remains to be seen in future studies whether the observed power comodulation generalizes to other muscles, including hand muscles.

### Spatial and Frequency Origin of the Leaking Oscillations

CST bandpower was particularly informative about the mental task around the IMC peak frequency. Furthermore, the bandpower correlations between CST and EEG were largest around the IMC peak frequency. The IMC reflects the strength of the common synaptic input, and the common synaptic input is at least partly of supraspinal origin (Farmer, 1998; Negro and Farina, 2011; Farina et al., 2014; Negro et al., 2016; Dideriksen et al., 2018). Thus, we observed the strongest correlation with cortical oscillations at a frequency where the common synaptic input, and so the cortical input, is believed to be high. Furthermore, the SPoC patterns suggest a centrally located dipole as a source, potentially in the foot area of the primary motor or somatosensory cortex. This finding is consistent with the somatotopic arrangement of CMC (Conway et al., 1995; Salenius et al., 1997; Chen et al., 2013; Mehrkanoon et al., 2014). Both findings indicate that CST oscillations modulated by mental tasks originate from the cortex, supporting the leaking-oscillation hypothesis.

The EEG SNR profile did not resemble the CST SNR profile. The EEG SNR was highest around the mu band, whereas the CST had low SNR in the mu band. This is not surprising as the cortical mu band is particularly modulated by motor imagery tasks (Pfurtscheller and Lopes da Silva, 1999), but supraspinal signals in the mu band are suppressed at the MN level (Williams and Baker, 2009; Williams et al., 2010).

For the mental tasks we studied, the discriminability of the EEG signal extended over the mu, beta, and low-gamma bands. However, the motor cortex oscillations modulated by these mental tasks were not transmitted downstream in a broadband manner, but preferentially around the IMC peak frequency.

### Using CST bandpower as a Control Signal for Neural Interfacing Applications

We were able to decode the performed mental task from CST features significantly better than by chance. However, the achieved classification accuracies were on average 30% (at a chance level of 25%). A significant improvement in classification performance is necessary to derive a useful control signal from task-dependent CST oscillations. We used a shrinkage LDA classifier which can offer close to state-of-the-art results in comparable BCI experiments with oscillatory cortical signals obtained by EEG (Lotte et al., 2018). Other classifiers, such as support vector machines or artificial neural networks, may improve performance, but the expected performance gain will probably be moderate (Schirrmeister et al., 2017; Lotte et al., 2018). A more promising approach involves intensive user training, as demonstrated in studies on brain-computer interfaces (Wolpaw and Mcfarland, 2004; McFarland et al., 2010; Collinger et al., 2013; Wodlinger et al., 2015). Initially, users would employ mental tasks as a control strategy to modulate CST bandpower. They would receive feedback on CST bandpower and learn to adjust it more accurately over the course of several weeks. In (Bräcklein et al., 2021), neurofeedback-based training using CST beta band power has been tested for a single session. Participants received visual feedback of their beta band power but were not told any explicit mental strategy on how to alter band power. Combining such feedback training with an initial mental strategy based on explicit mental tasks could help acquire control skills more quickly.

Moreover, non-invasive neural interfaces, such as EEG, may be combined with HD-EMG to derive more robust oscillation-based control signals, especially for able-bodied users. Future research should explore whether the discriminative information in CST signals about mental tasks is complementary or redundant to the information in non-invasively recorded brain signals.

### Influence of Task-Dependent Force Output

Beta oscillations do not directly translate into changes in muscle force (Farina et al., 2014). However, the opposite is not necessarily true, and task-dependent force output or dual-task interferences (Salmon and Thomson, 2007), e.g., due to shared mental resources, could confound the common bandpower changes in CST and EEG.

We found significant differences between mental tasks in force at the group level, and it was possible to classify tasks based on force features. Notably, the classification accuracies yielded by force features and CST bandpower features were uncorrelated across participants. Thus, task-dependent force changes are not a straightforward explanation for the classification results obtained with CST bandpower features. In contrast to force, the CST rate was not predictive of the mental task. This is surprising because the CST rate reflects the neural drive to a muscle and the force it produces. One possible reason for this is that MN decompositions over-represent MNs with higher recruitment thresholds, leading to an inaccurate estimation of the neural drive to the muscle (Caillet et al., 2024). In addition, at low forces, as in this experiment, changes in force are mainly implemented by changes in MN recruitment rather than changes in MN rates (Enoka and Duchateau, 2017). The CST rate may have remained largely unaltered by recruitment changes, as we only took into account a fixed set of predominantly active MNs when calculating the CST rate.

We removed the potential effect of force and CST rate on CST bandpower using multiple linear regression and only considering residuals for the analyses. We would like to note that only the effect of measurable force changes could be eliminated. Force changes along axes not captured by the force sensor could not be eliminated. Additionally, force changes might have been masked by the unavoidable slack in the coupling of the foot to the force sensor with a strap. However, removing the potential effect of force and CST rate had no impact on the classification accuracy with CST bandpower features, indicating that CST bandpower was not significantly affected by force or CST rate in the first place.

We found CST beta power changes to be in the same direction as EEG beta power changes (positive correlation). Task-dependent force changes do not explain this positive correlation as EEG beta power is not sensitive to force changes during a movement (Mima et al., 1999; Stancak et al., 1997; Zaepffel et al., 2013). Moreover, the correlations are not solely driven by the CST and EEG band power differences between the single-task condition (*no task*) and any dual-task condition (*foot MI, hand MI*, or *math* task) because the exclusion of *no task* still yielded significant CST-EEG correlations. Possible differences in task difficulty between the dual-tasks also do not explain the correlations as EEG beta and gamma bands are not sensitive to task difficulty in a dual-task experiment (Shaw et al., 2018). Most importantly, the task correlations between EEG and force-related parameters were, on average, close to zero, even though we used a spatial filter optimized to extract the force-related parameters. In line with this, potential force-related confounders had a negligible effect on the CST classification accuracies.

In conclusion, although we observed task-dependent force changes, the positive power correlations between CST and EEG in the beta band are unlikely to be due to these force changes to a significant extent.

### Limitations

We do not know if the coherence in the beta band (Conway et al., 1995; Halliday et al., 1998; Farmer, 1998; Marsden et al., 2000; Omlor et al., 2007; Gwin and Ferris, 2012; Ibáñez et al., 2021) is at least in part the result of leaking oscillations or a requirement for oscillations to leak downstream. In the case of the latter, oscillations may not be able to perpetually leak down as the coherence is decreased during dynamic movements (Feige et al., 2000; Kilner et al., 2003; Omlor et al., 2007; Mehrkanoon et al., 2014). This would also limit the applicability of CST bandpower as a control signal for neural interface applications, for example movement augmentation.

The information flow between the cortex and MNs in the beta band was found to be bidirectional by Witham et al. (Witham et al., 2011). Our analysis approach based on power correlation is unable to determine the directionality of information flow. While it is possible that some oscillations might be transmitted back to the cortex, this particular question is beyond the scope of this paper. Furthermore, our methodology cannot exlcude that the found correlations between EEG and CST result from a common input from a third, possibly subcortical source.

Recent research suggests that sensorimotor beta that appears on trial averages as a sustained oscillation may be, in fact, beta bursts on the single-trial level (Sherman et al., 2016; Barone and Rossiter, 2021). Beta bursts at the cortical and MN level were found to be time-locked (Echeverria-Altuna et al., 2022; Bräcklein et al., 2022) and were similarly altered in a neurofeedback scenario (Bräcklein et al., 2022). Our analyses are based on the variance of the EEG and CST within the task phase. The variance reflects the power within this period and is affected monotonically by the amplitude of sustained oscillations or, in the case of bursts, monotonically by their rate, duration, or amplitude. Due to this monotonic relationship, the identified comodulation between CST and EEG aligns with the leaking-oscillation hypothesis for both sustained oscillations and bursts. However, in the case of bursts, we cannot exclude the possibility that different burst parameters were comodulated. For example, CST burst *duration* may have been comodulated with EEG burst *amplitude*. Such potential cross-parameter comodulation would imply an intermediate processing step, possibly implemented in spinal networks, although its functionality would be unclear.

## 4 Conclusion

The execution of mental tasks led to common power changes in macroscale brain signals and spinal motor neurons. Furthermore, these power changes were discriminable across mental tasks at the spinal motor neuron level. As the mental tasks were different from the executed motor task, our findings corroborate the hypothesis that motor-unrelated cortical oscillation can leak downstream to spinal motor neurons.

## 5 Methods

### 5.1 Participants

Fifteen participants were recruited, including 11 men and 4 women. The participants were between 21 and 47 years old (28.3 ± 6.1 years; mean ± std. dev.). Recruitment was independent of any previous experience with electrophysiology experiments and handedness. The study was approved by the ethics committee of the University of Freiburg (approval number 21-1388-1), and written informed consent was obtained from the participants.

### 5.2 Experimental Paradigm

The participants were seated on a comfortable chair with their arms placed on armrests. Their right leg was fastened to a foot ankle dynamometer (NEG1, OT Bioelettronica SRL, Italy). A computer screen placed in front provided force feedback and instructions to the participants. We used a self-developed software package for this purpose (YAGA, https://yaga.readthedocs.io). We asked participants to perform an isometric maximum voluntary contraction (MVC) with their right foot (dorsiflexion). This was done to allow the presentation of the relative force value later in the recording session. Participants performed three MVCs for 5 s each. The maximum force value among the three contractions was selected as the MVC force value. The participants were then familiarized with the force feedback by tracking a trapezoidal force profile with their right foot five times (10 s ramp up, 5s hold at 10% MVC, 10 s ramp down). They were able to see the relative dorsiflexion force on the computer screen along with the target force. After that, the participants tracked a similar trapezoidal force profile four times but with a hold phase of 30 s.

Next, we instructed participants to perform a static isometric dorsiflexion with their right foot at 8% MVC while simultaneously performing one of four mental tasks, see Figure 1a. The four mental tasks were: motor imagery (MI) of right foot dorsi/plantarflexion (*foot MI* task), motor imagery of both hands opening/closing (*hand MI* task), continuously subtracting the number 3 from a random number displayed on the computer screen (*math* task), and no additional task (*no task* task). For the two MI tasks, the participants were instructed to perform kinesthetic MI (Neuper et al., 2005) of repetitive 0.5 Hz movements. The detailed sequence of a *trial* is shown in Fig. 1b. The participants started the isometric dorsiflexion at trial start and maintained the target force until the end of a trial. During the trial, the force feedback from the isometric dorsiflexion was provided by a ball moving on the computer screen’s vertical axis. At 5 s, the task cue was presented, indicating the mental task to perform. Participants performed the requested task for 5 s until the end of the trial. No feedback was given on the execution of the mental tasks. The time interval from 6 s to 9.5 s is referred to as the *task phase*. A *run* comprised 24 consecutive trials with a random inter-trial interval of 4 to 6 s. We recorded 15 runs with breaks of a least 1 min between runs, with the first run being a familiarization run. In total we recorded effectively 336 trials, i.e., 84 trials per task.

In the end, we again asked the participants to track four trapezoidal force profiles with a 30 s hold phase (data from trapezoidal force tracking tasks were not further used in this manuscript).

### 5.3 Recording

We recorded electrical activity from the tibialis anterior muscle using an HD-EMG grid electrode (GR08MM1305, OT Bioelettronica SRL, Italy). The grid contained 64 monopolar electrodes with 13 mm inter-electrode distance and was placed on the muscle belly aligned with the fibre direction. EMG reference and EMG ground electrodes were placed on the right and left ankle, respectively. Furthermore, we recorded bipolar EMG signals from the right gastrocnemius muscle and extensor digitorum and flexor digitorum muscles of both arms. The force signal was measured with a foot-ankle dynamometer connected to a pre-amplifier (forza, OT Bioelettronica SRL, Italy). The EMG and force signals were recorded using a biosignal amplifier (Quattrocento, OT Bioelettronica SRL, Italy) with a sampling rate of 2048 Hz. The EMG signals were filtered using a 0.7 to 900 Hz hardware bandpass filter. We recorded electrical brain activity with a 61 EEG channel montage of active Ag/AgCl electrodes covering frontal, central parietal and temporal areas. The EEG ground electrode was placed on Fpz; a dedicated physical reference electrode was not employed, i.e., the acquisition was reference-free. Additionally, we recorded electrooculography (EOG) signals with three electrodes placed above the nasion and below the outer canthi of both eyes. The EEG and EOG signals were recorded with a second biosignal amplifier (actiCHamp, Brain Products GmbH, Germany) and sampled at 2000 Hz with a 530 Hz anti-aliasing filter. The signals from both biosignal amplifiers were recorded and synchronised using Lab Streaming Layer (LSL) (https://github.com/sccn/labstreaminglayer). To access the data from the Quattrocento biosignal amplifier, we developed a custom LSL application (https://github.com/neurofreiburg/QuattrocentoLSLApp).

### 5.4 Preprocessing

The signals were processed with Matlab R2023b (MathWorks, USA) and the external toolboxes BioSig 3.7.6 (Schlogl and Brunner, 2008) and EEGLAB v2022.0 (Delorme and Makeig, 2004). To compute the relative force, we filtered the force signal using a median filter with a window length of 31 samples and normalized it to the participants’ MVC force levels. We referenced the EEG and EOG signals to AFz. Next, the (HD-)EMG, EEG, EOG and force signals were downsampled using a common time base to 1024 Hz for easier processing in terms of memory consumption and computation time. A 400 Hz anti-aliasing filter was applied before downsampling (4^th^-order zero-phase Butterworth filter). To suppress line interference, we applied a notch filter at 50 Hz and at the first two harmonics to the (HD-)EMG, EEG and EOG signals. Finally, we filtered the EEG/EOG data from 2 Hz to 150 Hz with a band-pass filter, and the force signal with a 10 Hz low-pass filter (4^th^-order Butterworth) to suppress non-physiological signal components.

We visually inspected HD-EMG, EEG and EOG signals and excluded noisy or bad channels. Possible stationary artefacts in EEG signals were removed with an independent component analysis (ICA) (Lee et al., 1999; Delorme and Makeig, 2004). We visually inspected ICs and removed components representing typical artefacts such as eye blinks, eye movements or muscle activity. Trials with EEG values exceeding ± 250 μV were excluded from the ICA and the inspection of ICs.

Finally, we excluded trials with transient artefacts in the *task phase* from the subsequent analyses. For the analyses relying solely on HD-EMG signals, we excluded trials with relative force values below 6% or above 10% MVC. Next, we excluded trials with abnormal force mean values or force standard deviations in the task phase. Moreover, we excluded trials with abnormal task-phase HD-EMG power values (wrt. the maximum across channels). Abnormal trials were defined as trials in which the respective statistic (i.e., force mean values, force std. dev., and HD-EMG power) deviated more than 7.413 times the median absolute deviation (MAD) from the median over trials. MAD is a robust alternative to the standard deviation and the threshold corresponds to 5 times the standard deviation for normally distributed data (Leys et al., 2013). In total, this led to an exclusion of 0% to 29% of the trials based on force and HD-EMG signals (6% ± 8%; mean ± std. dev. over participants). For the analyses relying solely on EEG data, we excluded trials with EEG amplitude values outside ± 75 μV, or abnormal task-phase EEG power values (wrt. the maximum across channels) using a threshold of 7.413 times the MAD. Furthermore, we computed joint probability and Kurtosis of the EEG amplitude values for each trial and channel using EEGlab (Delorme and Makeig, 2004). We excluded trials where either statistic exceeded a threshold of 5 times the standard deviation from the mean across trials for that statistic for any channel. In total, this led to an exclusion of 3% to 11% of the trials based on EEG signals (6% ± 3%; mean ± std. dev.). All criteria were combined for analyses relying on HD-EMG and EEG signals, resulting in the exclusion of 5% to 32% of the trials (12% ± 7%; mean ± std. dev.).

### 5.5 EMG Signal Decomposition and Cumulative Spike Train

A validated MU decoder from Negro et al. (Negro et al., 2016) was used to extract spiking activity of MNs from HD-EMG signals. The MN decoder was trained with HD-EMG data from all trials within the time interval of 5 s to 9.5 s relative to trial start. This decoder uses a blind source separation approach to decompose an HD-EMG signal into spike trains of MNs. To do this, the decoder algorithm extends the multi-channel HD-EMG signal with additional channels comprising time-lagged versions of the original HD-EMG signal, whitens the extended signal, and finds sparse source signals with a fixed-point algorithm (Hyvärinen, 1999; Hyvärinen and Oja, 2000). Typically, for signals sampled at around 2 kHz, the signal is extended with 16 successively increasing time-lags. Since we downsampled the signals to 1024 Hz, we selected 8 time lags to effectively cover the same time period. The fixed-point algorithm is followed by an iterative refinement step that uses a clustering approach to separate spikes from noise. A source signal was eventually accepted if it had a silhouette measure (SIL) (Negro et al., 2016) higher than 0.85. Due to the extension with time-lags, the decoder algorithm can find duplicate spike trains which are generated by a single MN but shifted by a constant time (the same spikes are detected at different time-lags). To detect duplicate spike trains, we applied agglomerative hierarchical clustering. We considered spike trains to be in the same cluster, i.e., belonging to the same MN, if they shared more than 5% of identical spikes. Here, we allowed one spike train to be shifted up to ± 32 samples relative to another to compensate for a possible constant time shift between spike trains. After compensating for this potential time shift, we considered spikes from two spike trains to be identical if they differed by a maximum of one sample.

We kept then only MNs (1) which had an average spike rate of more than 5 Hz, (2) where the standard deviation of the average firing rate across trials was less than 1.5 Hz (after detrending the spike rate with a moving average over 50 trials), and (3) where the trial-averaged coefficient of variation (CoV) of the inter-spike interval was less than 0.5. We considered only the task phase for all three criteria. This yielded MN counts between 2 and 35 (15.9 ± 10.5; mean ± std. dev. over participants).

The CST was eventually obtained by summing the individual spike trains of the MNs.

### 5.6 Power Comodulation Analysis

We analysed whether oscillations in the CST were comodulated with brain signals (see Figure 1c). To do this, SPoC (Dähne et al., 2014) was used to find a linear projection of EEG channels that maximizes covariance with CST in bandpower. In other words, SPoC found a spatial filter that extracted an EEG source, whose bandpower had maximal covariance with the CST bandpower.

First, we applied a filter bank from 5 to 74 Hz to the CST. The filter bank consisted of 5 Hz wide bands, separated by 1 Hz steps (i.e., 5-9, 6-10, …, 70-74 Hz; 4^th^-order zero-phase Butterworth). We calculated the variance over the task phase to obtain the band power *CSTBP*_*f,i*_:

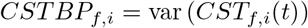

where *CST*_*f,i*_(*t*) is the *i*^*th*^ trial of the bandpass filtered CST of frequency band *f*, and *t* indexes the time within the task phase of the trial. Subsequently, the signals were detrended to compensate for possible non-stationarities during a recording session. For that, we subtracted the moving median using a window length of 50 trials and obtained the detrended band power:

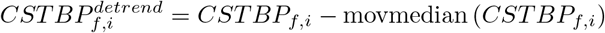

Potential *task-dependent* changes in force output could have confounded CST band power. To remove possible force-related confounders, we computed the mean value and standard deviation of the force as well as the CST rate during the task phase. We then trained for each frequency band a multiple linear regression model with force mean, force standard deviation, and CST rate as predictor variables, and 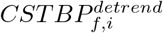 as response variable. Only the residuals 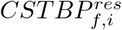 were then used for further analysis. The residuals corresponded to the original CST bandpower features but with the linear effects of force and CST spike rate removed. Next, we applied the same filter bank to the EEG signal and applied SPoC separately for each frequency band. The SPoC implementation provided in (Meinel et al., 2019) with shrinkage (Ledoit and Wolf, 2004; Schäfer and Strimmer, 2005) was used for that. We calculated the variance over the task phase of the EEG channel projection found by SPoC (only the first SPoC component was used):

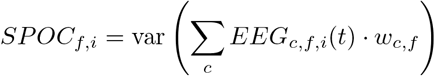

where *EEG*_*c,f,i*_ is the EEG signal with *c* indexing the EEG channel, and *w*_*c,f*_ the weight found by SPoC. SPoC found the weights *w*_.,*f*_ such that the covariance over trials between *SPOC*_*f,i*_ and 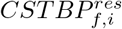 was maximized. Importantly, *SPOC*_*f,i*_ was found using a 10-fold cross-validation to prevent overfitting of the SPoC weights. Thus, the computation of the SPoC weights and their application was performed on separate trial sets obtained by cross-validation. Subsequently, we calculated a robust average of CST and SPoC trials over the mental tasks:

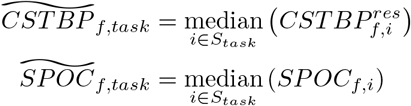

where *S*_*task*_ is the set of trial indices belonging to *task*.

Eventually, the correlation over individual trials (*trial* correlation) and over tasks (*task* correlation) was computed. To this end, we computed the *Kendall rank correlation coefficient τ* between CST and SPoC variances for each frequency band to identify comodulation. *τ* is a robust correlation measure as it is based on rank correlation. The *trial* correlation was calculated after removing the effect of the task from CST and SPoC trials:

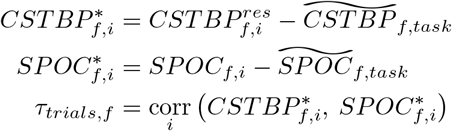

where *task* is the performed task in trial *i* and *corr* refers to the Kendall rank correlation coefficient. The *task* correlation was computed as the correlation over task averages:

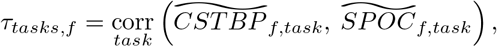

It should be noted that *τ*_*tasks,f*_ was computed using only four data points (mental tasks) per participant. This limited sample size resulted in a high variance of *τ*_*tasks,f*_, which was addressed with statistical tests conducted across participants. We computed *τ*_*tasks,f*_ using the Matlab implementation of Kendall’s *τ* coefficient, which includes an adjustment for ties, known as Kendall’s *τ*_*b*_ (Gibbons and Chakraborti, 2003). *τ*_*b*_ is identical to the originally proposed Kendall’s *τ* coefficient (*τ*_*a*_) in the absence of ties in any variable. *τ*_*a*_ is known to be an *unbiased* estimator for any bivariate distribution (Gibbons and Chakraborti, 2003). Furthermore, for independent continuous random variables, *E*[*τ*_*a*_] = 0 (Gibbons and Chakraborti, 2003). Since CST or SPoC trials did not have any ties, i.e., no CST or SPoC trials had exactly the same power values, our estimation of Kendall’s *τ* coefficient was unbiased. Thus, the small sample size did not introduce any systematic error.

The trial and task correlation coefficients were computed for each participant and frequency band. We performed group-level analyses on (1) the correlation coefficients of individual frequency bands and on (2) the correlation coefficients averaged across frequency bands covering 15 to 45 Hz (*band-averaged* correlations). This frequency range was chosen to cover the beta and low-gamma band of potentially transmitted brain oscillations. Furthermore, (3) we averaged for each participant the correlation coefficients of the 5 neighbouring frequency bands centred at the frequency with the highest common synaptic input, as determined by intramuscular coherence (*IMC-aligned* correlations). For the calculation of the IMC, the MNs were first randomly split into two equally sized sets. We computed the CSTs of both sets (Bräcklein et al., 2021) and applied a short-time Fourier transform (STFT) to the set-CSTs using a 1 s Hamming window. Next, we computed the power spectra of both set-CSTs and their cross-power spectrum, averaged them over the task phases, and calculated the coherence (Bastos and Schoffelen, 2016) between both MN sets. The random MN splitting was repeated in total 500 times, and the coherences were eventually averaged over all random splits to obtain the IMC. We determined the peak frequency of the IMC between 15 and 45 Hz in order to find the frequency with the largest common synaptic input. The peak frequency was defined here as the frequency of the peak with the largest prominence (Kirmse and de Ferranti, 2017).

### 5.7 Discriminability Analysis

We analysed whether the mental tasks modulated the power of the CST oscillations. To this end, the discriminability of the CST band-power 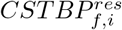 was evaluated with SNR analysis and single-trial classification.

#### SNR Analysis

The SNR at each frequency band during the task phase was quantified. For that, we defined the signal as *task-dependent band-power variability* (i.e., variance of task-averaged CST band-power):

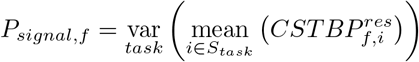

where *S*_*task*_ is the set of trial indices belonging to *task*. We defined the noise as *task-independent band-power variability* (i.e., variance of CST band-power across trials after removing the average task band-power):

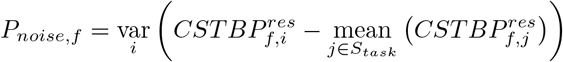

where *task* is the performed task in trial *i*. As the variance operator involves calculating the mean, we used the mean rather than the median to determine the task-averaged CST band power values for consistency. We did not use MAD as an alternative to the variance in the calculation of the SNR due to the limited number of data points (i.e., 4 tasks). The SNR was eventually calculated as:

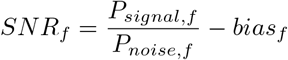

where the bias term *bias*_*f*_ was determined with non-parametric bootstrap sampling (Efron and Tibshirani, 1994) using 5000 repetitions. We computed the SNR for the frequency bands 5-9, 6-10, …, 70-74 Hz. Additionally, we calculated the average SNR over the frequency range of 15 to 45 Hz (*band-averaged* SNR), and the average SNR over the 5 adjacent frequency bands centred at the IMC peak (*IMC-aligned* SNR). A baseline SNR *SNR*_*baseline,f*_ was computed using the same procedure as described before, but using the time interval 2.5 to 5 s relative to the trial start where no task information was available yet. Furthermore, we computed the SNR using the central EEG electrode Cz. For this analysis, a Laplace spatial filter was applied to extract brain activity predominantly from the motor cortex (Mcfarland et al., 1997; Tsuchimoto et al., 2021).

#### Single-trial Classification

We quantified the single-trial discriminability of CST band power features (*CST-FB*) with an sLDA classifier (Peck and Van Ness, 1982; Blankertz et al., 2011). The shrinkage parameter was computed analytically using the method of Ledoit and Wolf (Ledoit and Wolf, 2004; Schäfer and Strimmer, 2005). The features were the band power values 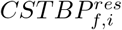 computed with a filter bank as in Section 5.6 but with a frequency range covering beta and low gamma bands in 2 Hz steps (i.e., 15-19, 17-21, …, 41-45 Hz). Additionally, we classified CST band power features (*CST-IMC*) obtained only from the participant-specific IMC peaks and the 4 neighbouring frequency bands (stepsize 1 Hz). Furthermore, we evaluated the discriminability of EEG signals and extracted band power features from all 61 monopolar EEG channels using the same filter bank as the CST-FB features. We were only interested in EEG frequencies which could potentially leak down to MNs. We therefore omitted the often task-discriminative mu-band as signals around 10 Hz are attenuated when propagating downstream (Farina and Negro, 2015). We also quantified the discriminability of potential confounders. This was done by calculating the mean and standard deviation of the force, as well as the CST rate, within the task phase (*FORCE-MEAN, FORCE-STD* and *CST-RATE*). We then computed the classification accuracies yielded by the features using an sLDA classifier and a 10 × 10 fold cross-validation.

The agreement of classifier predictions from CST-FB and EEG-FB features was evaluated with the *uncertainty coefficient* (Press et al., 2007). *The uncertainty coefficient U* is a measure of nominal association derived from information entropy ranging between 0 and 1:

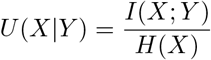

with *I*(*X*; *Y*) being the mutual information between *X* and *Y*, and *H*(*X*) the entropy of *X. U* (*X Y*) is not symmetric with respect to *X* and *Y* and a symmetric uncertainty coefficient can be obtained as (Press et al., 2007):

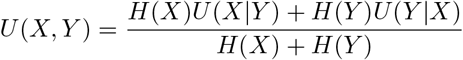

For each participant, we first computed the task predictions separately from CST-FB and EEG-FB features using a 10 × 10 fold cross-validation with common train and testsets. Subsequently, we computed *U* (*X, Y*) across all testsets of a cross-validation repetition (i.e., a reshuffling of trials). We then removed the bias with non-parametric bootstrap sampling (Efron and Tibshirani, 1994) using 1000 repetitions, and eventually averaged *U* (*X, Y*) over all 10 cross-validation repetitions.

### 5.8 Statistics

Non-parametric tests were used when comparing one or two samples. For these comparisons, except when involving *U* (*X, Y*), we used a two-sided Wilcoxon signed-rank test. The Wilcoxon signed-rank test tests the null hypothesis that a sample or the difference between two paired samples comes from a symmetric distribution with a median of zero. The significance level was set to *α* = 0.05. For multiple comparisons, the false discovery rate was controlled at *q* = 0.05 with the Benjamini & Hochberg procedure (Benjamini and Hochberg, 1995), and we report Benjamini & Hochberg adjusted p-values (*p*_*adj*_) (Yekutieli and Benjamini, 1999). Adjusted p-values reported in the same paragraph in Section Results were considered as a set to which the Benjamini & Hochberg method was applied. We evaluated the statistical significance of the symmetric uncertainty coefficient *U* (*X, Y*) on the participant-level with a one-sided permutation test (Nichols and Holmes, 2001; Maris and Oostenveld, 2007), where the CST-FB and EEG-FB labels were permuted across all testsets within a cross-validation repetition. We sampled *N* = 10000 symmetric uncertainty coefficients from the permutation distribution. The p-values were calculated as *p* = (*b* + 1)*/*(*N* + 1), where *b* is the number of symmetric uncertainty coefficients from the permutation distribution greater than or equal to the observed symmetric uncertainty coefficient (Phipson and Smyth, 2010). A one-sided test is justified as any value of *U* (*X, Y*) smaller than the permutation distribution has to be considered as a random fluctuation. We used repeated measures ANOVA for comparisons involving more than 2 samples. If the sphericity assumption was violated (Mauchly’s test), a Greenhouse-Geisser (GG) correction was applied. Post-hoc analysis was performed with a Tukey-Kramer test. Confidence intervals of statistics based on one or two samples were computed at a level of 95% using bootstrapping with the bias-corrected and accelerated percentile method (DiCiccio and Efron, 1996).

## Acknowledgements

The authors thank Meng-Jung Lee for discussions and critical feedback.

## Author Contributions

P.O., D.F., C.M. designed research; P.O. acquired and analysed data; P.O., D.F., C.M. interpreted data; P.O. wrote the initial draft; P.O., D.F., C.M. reviewed and edited the paper.

